# Retrograde transduction of dopaminergic cells in substantia nigra of rhesus monkey

**DOI:** 10.64898/2026.01.07.697726

**Authors:** Anya S. Plotnikova, Walter Lerchner, Alexander C. Cummins, Gang Chen, Leonardo Salhani, Vincent D. Costa, Bruno B. Averbeck, Barry J. Richmond, Zayd M. Khaliq, Mark A. G. Eldridge

## Abstract

Neuromodulatory systems regulate neural circuits across broad regions of the brain, and disruption of the dopaminergic system contributes to psychiatric and neurodegenerative disorders. Engineered viral vectors have been used to target the neuromodulatory systems of the nonhuman primate brain (Y. Chen et al., 2023; El-Shamayleh et al., 2016; Gray et al., 2010; Lerchner et al., 2014; Perez et al., 2022). However, a conspicuous obstacle to the isolation and modulation of specific pathways is the inability of many retrogradely infecting viruses to transduce dopaminergic (DA) cells efficiently (Tervo et al., 2016; Cushnie et al., 2020; Weiss et al., 2020). We compare the DA neuron retrograde transduction efficacy of four viral vectors after injection into the striatum of nonhuman primates (NHP). Selectivity was assessed by comparing the neuronal co-expression of fluorescent reporter protein and tyrosine-hydroxylase (TH) antibody in substantia nigra pars compacta (SNc). The rabies pseudotyped lentiviral vector, FuG-B2, produced superior retrograde transduction of DA cells to FuG-C or FuG-E. AAV2.retro was the least effective.

## 1 Introduction

Dopamine signaling dysfunction is implicated in many psychiatric and neurodegenerative conditions (H. Chen et al., 2024) including substance use disorder (Volkow et al., 2009), Parkinson’s disease (Damier et al., 1999; Fearnley & Lees, 1991; German et al., 1992), and dystonia (Karimi & Perlmutter, 2015; Ribot et al., 2019). The substantia nigra (SN) is the primary source of dopaminergic (DA) innervation in the mammalian central nervous system (CNS) (Björklund & Dunnett, 2007; Joel & Weiner, 2000). The SN is anatomically and functionally divided into two distinct regions: the substantia nigra pars compacta (SNc), which contains the majority of DA neurons projecting to the striatum, and the substantia nigra pars reticulata (SNr), which consists primarily of GABAergic neurons (Kelly et al., 2022).

Nigrostriatal DA neurons exhibit functional diversity which underlies their essential roles in motor control and goal-directed behaviors (Freeze et al., 2013; Howe & Dombeck, 2016). As critical components of the reward circuit, this population of DA neurons contributes to motivation, aversion, and learning processes (Bromberg-Martin et al., 2010). Selective modulation of specific sub-populations of DA cells via chemogenetic or other gene therapy methods holds great potential for clinical applications and for basic research on the neural circuitry governing DA-dependent processes (Haber & Knutson, 2010). Targeting the somata of dopaminergic neurons, however, poses a challenge due to their relatively inaccessible location in the ventral midbrain. Consequently, local injection results in non-selective targeting of all DA neurons within the region, thus requiring a 2-vector system for pathway-specific manipulations (Isa, 2022; Koshimizu et al., 2021; Vancraeyenest et al., 2020). An alternative approach is to inject a larger, more accessible brain region, such as the striatum, to retrogradely infect dopamine terminals. Retrograde targeting would allow for selective manipulation of specific dopaminergic pathways while preserving the functionality of other DA projections (Yan et al., 2025). In conjunction with retrograde targeting, cell-type specific regulatory elements, and vectors with selective tropism can constrain expression to DA neurons of interest (Blömer et al., 1997; Cockrell & Kafri, 2007; Naldini et al., 1996).

Engineering vector tropism allows for high efficacy and specificity in the transduction of target cells (Chan et al., 2017; Eldridge & Galvan, 2023). Viruses that have been engineered to confer enhanced retrograde transduction properties include a variant of AAV2, AAV2.retro (Tervo et al., 2016), and HIV-1-based lentiviral vectors (Carpentier et al., 2012) pseudotyped with rabies Fusion Glycoproteins B, C, and E (FuG-B2, FuG-C, and FuG-E, respectively) (Cronin et al., 2005; Kato & Kobayashi, 2020). These engineered viral vectors are endowed with the ability to be internalized by most axon terminals and transported to neuronal cell bodies. HIV-1-based retrograde lentiviral vector, FuG-B2 (‘HiRet’), transduces all brain cell types (glia and neurons), whereas FuG-C (‘NeuRet’) and FuG-E are neuron specific (Kato & Kobayashi, 2020; Tanabe et al., 2019). The rapid propagation and toxicity of conventional retrograde viral techniques, such as rabies and herpes simplex, makes pseudotyped lentivirus and AAV more appealing vectors for long-term studies given their lower cytotoxicity and inflammatory potential (Tanabe et al., 2019; S.-H. Chen et al., 2023; Cushnie et al., 2020; Liu et al., 2022). Efficiency of retrograde transduction is dependent on vector, species, and brain region, among other factors. The AAV2.retro vector, though successful in transducing neurons in cortical and some subcortical structures of the mouse (Wang & Zhang, 2021), has shown inconsistencies in transduction efficacy of dopaminergic cells in the SN of both mouse (Tervo et al., 2016) and NHP (Albaugh et al., 2020; Cushnie et al., 2020; Weiss et al., 2020; Yan et al., 2025). Conversely, prior studies have shown that FuG-C and FuG-E efficiently transduce cells in the SNc (Kato, Kuramochi, et al., 2011; Kato et al., 2014). Though pseudotyped lentivirus can transduce DA neurons in the nonhuman primate, certain serotypes produce an inflammatory response when combined with the ubiquitous MSCV promoter (Kato, Kobayashi, et al., 2011; Tanabe et al., 2017, 2019). This study was designed to investigate the retrograde transduction efficiency and selectivity for dopaminergic cells of four different viral vectors, FuG-B2, FuG-C, FuG-E, and AAV2.retro.

## 2 Materials and Methods

### 2.1 Vector preparation and storage

Lentivirus was produced at titers of 10^9 I.U. using methods similar to those described previously (Lerchner et al., 2023). 293T (Lenti-X Invitrogen 632180, Life Technologies) cells were transfected with the Lenti-backbone plasmid and packaging plasmids (Addgene, Cambridge, MA, USA). The lentiviral vectors were pseudotyped with rabies virus glycoprotein variants (FuG-B2, FuG-C, and FuG-E) and engineered to express green or red fluorescent reporter proteins (GFP or RFP) under the human synapsin (hSyn) promoter which allows for neuron-specific expression. AAV2.retro-hSyn1-GCAMP6f-P2A-nls-dTomato was provided by Jonathan Ting (Addgene plasmid # 51085 ; http://n2t.net/addgene:51085 ; RRID:Addgene_51085) and AAV2.retro-hSyn-CRE-eGFP was received from James M. Wilson (Addgene plasmid # 105540 ; http://n2t.net/addgene:105540 ; RRID:Addgene_105540). (Table 1).

**Table 1.**
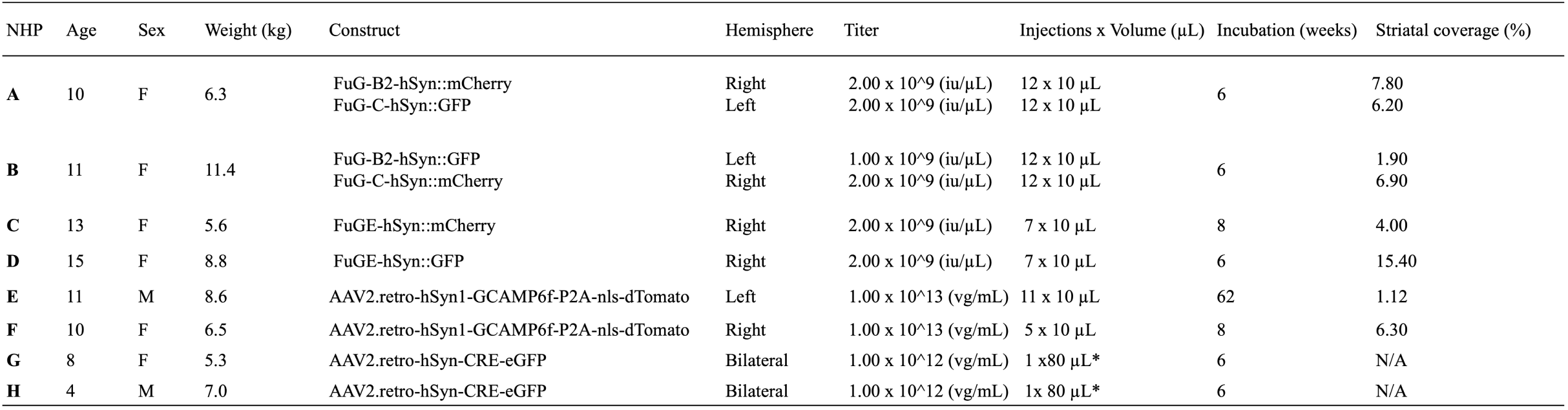
Summary table of surgical cases. Virus was targeted to striatum in stereotaxic surgeries. Incubation is defined as the time between viral injection and post-mortem fixation. Expression as a percentage in the rhesus macaque brain at the site of local injection averaged across 3 anterior-posterior levels. Asterisk (*) indicates use of convection enhanced delivery.

### 2.2 Subjects

All experimental procedures conformed to the Institute of Medicine Guide for the Care and Use of Laboratory Animals and were carried out under an Animal Study Proposal approved by the Animal Care and Use Committee of the National Institute of Mental Health. Subjects were 8 adult rhesus monkeys (*Macaca mulatta*: Monkey A, Monkey B, Monkey C, Monkey D, Monkey E, Monkey F, Monkey G, and Monkey H). All NHPs (6 females and 2 males) were between 4 and 15 years old, and their weights ranged from 5.3 to 11.4 kg (Table 1). They were housed on a 12-hour light-dark cycle with access to water ad libitum and fed a diet of monkey biscuits twice daily, approved treats, and fruit.

### 2.3 Surgeries

Surgeries were performed under aseptic conditions in a fully equipped operating suite under veterinary supervision. Prior to surgery, animals were sedated with ketamine hydrochloride (10 mg/kg, intramuscular) and midazolam (0.1 mg/kg, intramuscular). The NHPs were then intubated. A surgical level of anesthesia was induced and maintained with isoflurane (1–4%) throughout the procedures. Vital signs (body temperature, heart rate, blood pressure, SpO2, and expired CO_2_) were monitored. NHPs received antibiotic (Cefazolin) and isotonic fluids intravenously throughout the surgery. The brain was accessed by retracting muscle, removing a bone flap, and reflecting the dura mater. Stereotaxic injection coordinates were derived from pre-operative structural MRIs (Saunders et al., 1990; Walbridge et al., 2006). Post-operative MRI using manganese (Mn^2+^) contrast agent allowed for rapid post-operative visualization of viral vector delivery sites. The procedure has been described in detail previously (Fredericks et al., 2020). Postoperative visualization with Mn^2+^ suggested needle occlusion in NHP B during intrastriatal delivery of the FuG-B2 pseudotyped vector (see Fig. S1). Virus aliquots were stored at -80°C and were allowed to thaw on ice prior to injection. Virus was taken up into 100 μL Hamilton syringes (30G, beveled 45°, step) via a Harvard Apparatus Pump 11 Elite Nanomite.

For monkeys A – F, between 70 and 150 μL of virus (10 μL/site at a rate of 0.5 – 1.0 μL/min) was delivered unilaterally into striatum across three anterior-posterior levels (interaural +24 mm to +19 mm), 1.5 – 2.0 mm apart, of each NHP. Injections were matched across hemispheres in the FuG-B2 and FuG-C cases (Fig.1, Table 1). Injections along the same track were placed 2 –2.5 mm apart in the dorso-ventral plane. For monkeys G & H, single injections of AAV2.retro-hSyn-CRE-eGFP were delivered bilaterally into the ventral striatum, specifically targeting the nucleus accumbens (NAc). Using a progressive increase in infusion rate known as convection enhanced delivery (CED), 80 µL of virus was injected into each hemisphere at a rate of 0.5 µL/min to 3-3.5 µL/min over the course of 45 minutes. Methods for CED have been described previously in detail (Lee et al., 2025). To prevent infection and minimize swelling, NHPs A through F received dexamethasone sodium phosphate (0.4 mg/kg) and Cefazolin antibiotic (15 mg/kg) one day prior to surgery and in the week following the surgical date. Analgesic ketoprofen (10-15 mg) was given immediately following the surgery and for two days after. NHPs G and H animals were post-operatively monitored for 5–7 days by veterinary staff and received Cefazolin, Hydromorphone, and Buprenorphine (antibiotic and pain management).

**Figure 1.**
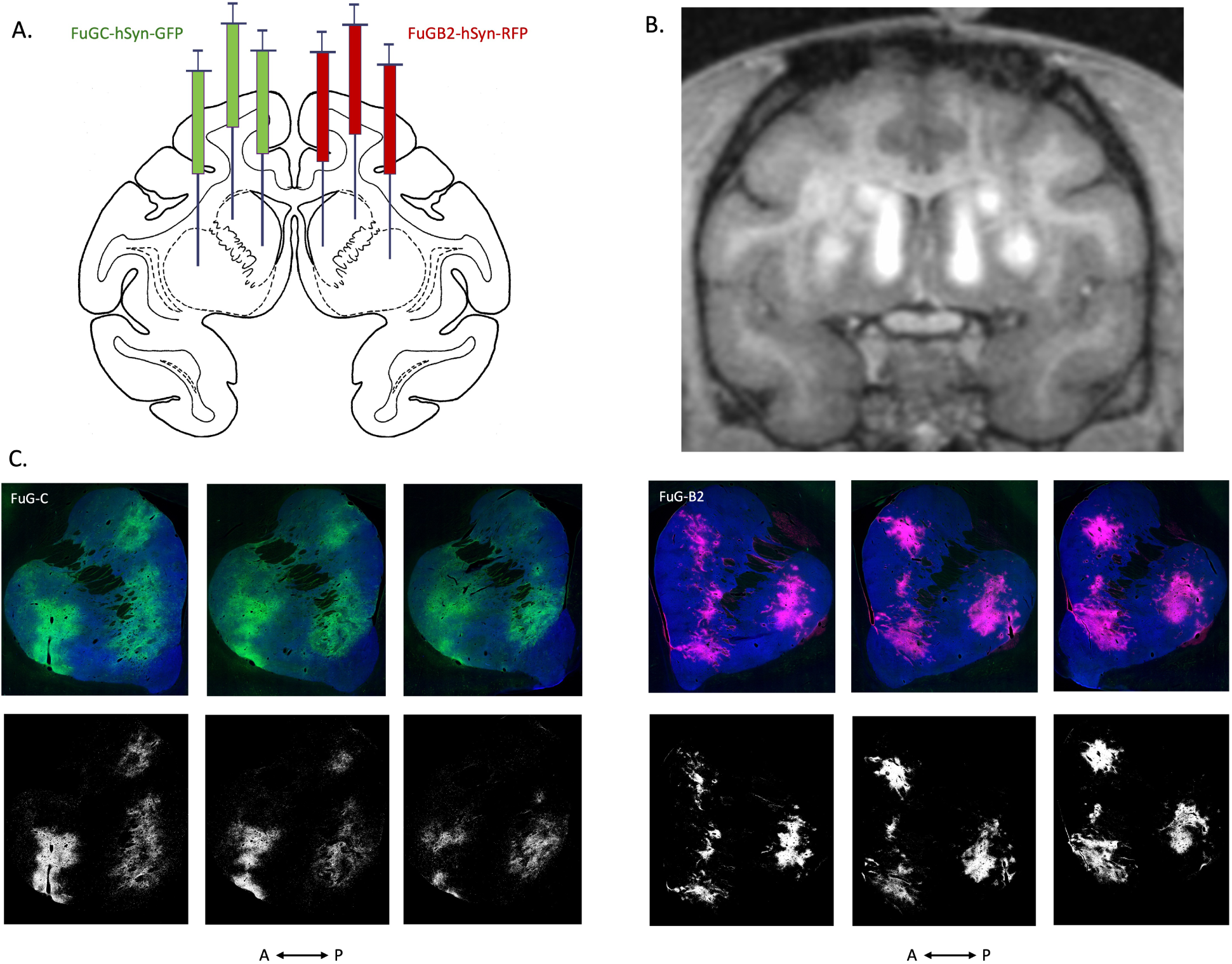
Viral delivery and expression in the striatum of a macaque brain. (A) Schematic illustrates striatal injection sites in Monkey A. FuG-C pseudotyped vector was delivered into the left hemisphere. FuG-B2 pseudotyped vector was delivered into the right hemisphere. (B) Same-day post-op MRI to visualize target accuracy via 0.2 mM manganese contrast agent co-infused with virus. (C) Reporter expression visualized post-mortem with immunohistochemistry and fluorescence microscopy across 3 consecutive anterior-posterior sections with 400 µm spacing. The image was thresholded using ‘intermodes’ on ImageJ (see methods) for quantification of the injection area (bottom row).

### 2.4 Tissue preparation and histological processing

After a minimum 6-week incubation period, the NHPs were euthanized. NHPs were administered ketamine hydrochloride (10 mg/kg), and were then intravenously given a lethal dose of sodium pentobarbital (100 mg/kg). Following euthanasia, the NHPs were transcardially perfused with 1 liter of saline and 3 liters of 4% paraformaldehyde in 0.1 M phosphate-buffered saline. Brains were extracted and refrigerated in 10% formalin. The brains were cryoprotected in buffered formalin/glycerol and then coronally sectioned with a freezing microtome into 40 µm coronal slices. Sections were collected in series of 10. Every tenth section was mounted on gelatin-coated slides, defatted, stained with thionin, and cover-slipped. Two series were stained with 3,3’-diaminobenzidine (DAB) for RFP and GFP, respectively. A series was processed for fluorescence immunohistochemistry (IHC) (with the exception of monkeys G & H, which were stained for DAB only). Sections were blocked in 5% normal donkey serum (vol/vol) and 0.3% Triton X-100 (vol/vol) in tris-buffered saline (TBS), incubated with rabbit anti-GFP primary antibody (1:10,000, Abcam AB290) in blocking buffer, washed and incubated with a biotinylated goat anti-rabbit IgG antibody (1:500, Vector Laboratories BA-1000). Signal was amplified with the VECTASTAIN ABC System (Vector Laboratories) according to specifications. For triple-fluorescence detection, NHP tissue sections were washed in PBS (3×10min), blocked with 20% goat serum, then washed again in PBS (3×10min) before overnight incubation at room temperature with primary antibodies: anti-GFP (Abcam AB13970, chicken, 1:5000), anti-mCherry (Abcam Ab167453, rabbit, 1:5000), and anti-TH (Immunostar 22941, mouse, 1:1000) diluted in PBS. Following primary incubation, sections were washed in PBS (3×10min), then incubated for 2 hours at room temperature with secondary antibodies: goat anti-chicken 488, goat anti-rabbit 555, and goat anti-mouse 647 (all Invitrogen, 1:300 in PBS), keeping sections in dark conditions to prevent photobleaching. After final PBS washes (3×10min), sections are mounted with DPX and coverslipped, resulting in triple-label fluorescence with GFP, mCherry, and TH signals. Technical issues during the processing of the AAV2.retro striatal sections resulted in compromised tissue quality, rendering the striatal dataset unsuitable for analysis.

### 2.5 Imaging and cell quantification

Mounted tissue sections were scanned with 10x (Fig. 1-3, and Fig. S1) and 40x (validation) objectives using an Olympus VS200 scanning microscope with fluorescence microscopy. Fluor 488 and Fluor 555 antibodies were visualized under a 3 ms exposure, and Fluor 647 was scanned under a 40 ms exposure. Brightness and contrast were matched across cases. Prior to manual quantification, cell identification criteria were determined based on morphology, luminance, size, and focal plane. The SNc boundary was determined using the cell distribution on a thionin-stained coronal section. The boundary was transposed onto a TH-stained fluorescent image of an adjacent series using anatomical landmarks and blood vessels. Boundaries were adjusted according to the dorsal/ventral distribution of TH-antibody positive cells. Using FIJI software (Schindelin et al., 2012), total SNc area of the imaged tissue was quantified in mm^2^. Individual reporter-positive cells and dopaminergic cells were quantified and plotted using Adobe Illustrator. Images were separated channel-wise and later overlaid to validate reporter and TH-antibody co-expression per cell. For quantification of expression at the injection site in striatum (interaural +24 mm to +19 mm), three consecutive coronal sections with the highest local expression were selected from each NHP and viral vector case. Exposure time, contrast, and brightness levels were matched across images. In FIJI, the striatum was outlined on a Fluor 647 section and quantified in square pixels. The injection area was visualized in the red or green channels (corresponding to the color of the reporter) with ‘Intermodes’ thresholding, and quantified in square pixels as a percentage of the total striatal area. Due to the absence of fluorescent striatal sections in the AAV2.retro cases, injection site quantification was performed on DAB sections rather than fluorescent images. A similar procedure and thresholding technique was used in FIJI to evaluate local expression at the striatal injection site. Ratios between the quantified injection site area and the total area of the striatum were used to compare the levels of viral expression at the site of local injection between cases (Table 2). SN and VTA in the AAV2.retro-hSyn-CRE-eGFP cases (NHP G and H) were scanned using 20x extended depth of field (EDF) with a 10 micron Z-stack on a DAB-stained section (Fig. S2).

**Table 2.** Viral vector expression and selectivity in SNc. SNc area, cell counts, and percent co-expression are averaged across 3 anterior-posterior sections per monkey for each viral vector case. Asterisk (*) indicative of convection enhanced delivery.

| Viral Vector | SNc Area (mm<sup>2</sup>) | Reporter<sup>+</sup> Cells | Reporter<sup>+</sup>/TH<sup>+</sup> Cells | Percent Co-expression |
| --- | --- | --- | --- | --- |
| **FuG-B2** (NHP A) | 7.9 | 269 | 260 | 97 |
| **FuG-B2** (NHP B) | 8.8 | 56 | 54 | 96 |
| **FuG-C** (NHP A) | 8.5 | 63 | 56 | 89 |
| **FuG-C** (NHP B) | 9.3 | 42 | 38 | 91 |
| **FuG-E** (NHP C) | 5.6 | 24 | 23 | 96 |
| **FuG-E** (NHP D) | 5.0 | 13 | 10 | 77 |
| **AAV.retro** (NHP E) | 7.1 | 2 | 1 | 50 |
| **AAV.retro** (NHP F) | 9.4 | 0 | 0 | 0 |

### 2.6 Statistical analysis

The data was fitted in a hierarchical model under the Bayesian framework. The model was formulated as below:

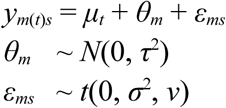

The input data *y_m_*_(*t*)*s*_ denotes the cell counts per unit area on slice *s* (*s* = 1, 2, 3) of the *m*th monkey (*m* = 1, 2, …, 6) that belongs to *t*th treatment group (*t* = 1, 2, 3, 4). The parameters *μ_t_*, *θ_m_*, and *ε_ms_* represent group-level effect, monkey-specific effect, and slice-level effect, respectively.

Noninformative hyperprior was adopted for *μ_t_*, weakly-informative prior (i.e., Student’s half-𝑡(3, 0, 1)) was used for standard deviations *τ* and *σ*. The hyperprior for the degrees of freedom, 𝜈, of the 𝑡-distribution for *ε_ms_* was Γ(2,0.1).

The above hierarchical model was numerically solved using the R (R Core Team, 2024) package *brms* (Bürkner, 2017). The resulting posterior samples were used to generate the six pairwise comparisons among the four treatment groups using the R package ggplot2 (Wickham, 2016).

## 3 Results

### 3.1 Efficiency of viral vector transduction of SNc DA neurons

We injected four viral vectors unilaterally into the striatum of six NHPs (n = 2 hemispheres per vector) and compared their ability to efficiently transduce cells in the SN of the rhesus monkey (Fig. 1, Table 1). Post-operative visualization of the injected region revealed uniform signal (via Mn^2+^ contrast agent) in the target areas, consistent with planned trajectories (Fig.1B), with the exception of the left hemisphere of NHP B (Fig S1B-C). Post-mortem fluorescent IHC of reporter expression was consistent with post-operative Mn^2+^ signal distribution (Fig. 1C).

Fluorescent microscopy confirmed successful retrograde transduction of neuronal populations following intrastriatal injection (Fig. 2A).

**Figure 2.**
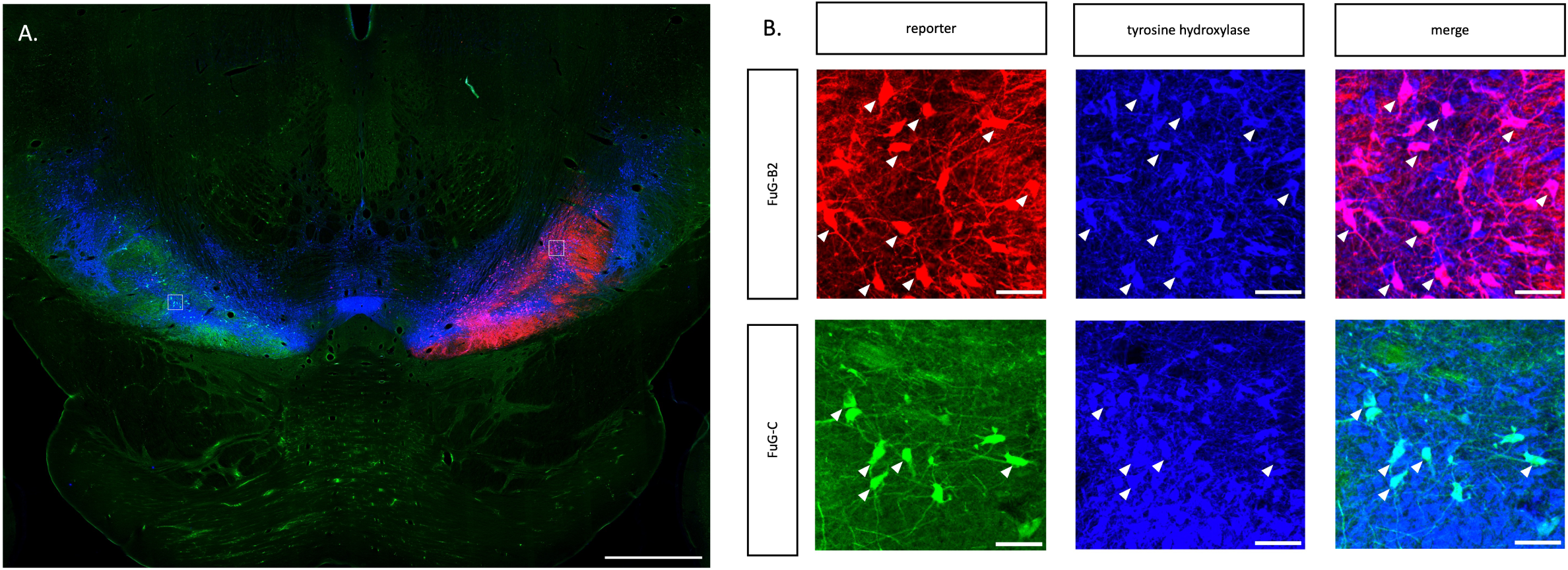
Co-expression of fluorescent reporter protein and TH antibody in the SN of a macaque visualized using immunohistochemistry and fluorescence microscopy. (A) Sample coronal section from Monkey A. Scale bar 2 mm. (B) High magnification images corresponding to the white boxes in A. Reporter and TH IHC in SN after transduction with FuG-B2 and FuG-C viral vectors. White arrows indicate individual retrogradely labeled cells co-expressing reporter protein and TH. Scale bar 100 µm.

Analysis of individual channels and merged images revealed co-expression of reporter and TH-antibody, suggesting that the neuronal populations transduced were dopaminergic. Striatal expression was not predictive of which vectors would show significant transduction, as in some cases substantial intrastriatal coverage did not yield robust expression of reporter in DA cells (Tables 2, 3).

Reporter expression in the SNc was quantified across three consecutive anterior-posterior sections within a series for each viral vector (Fig. 3, Table 2). The highest level of reporter expression in SNc was observed after intrastriatal delivery of FuG-B2 (‘Hi-Ret’) (Fig. 4A, Tables 1, 2). FuG-C and FuG-E produced comparable levels of transduction in SNc, though lower than FuG-B2 (Fig. 4A, 5, Table 2), despite similar striatal coverage (Table 1). AAV2.retro produced negligible expression in SNc (Fig. 3B, 4A, Table 2), despite expression in other brain regions, including the basolateral nucleus of the amygdala (BLA) (Fig. 6). The data was fitted to a hierarchical model under the Bayesian framework, accounting for variability between subjects and measuring the density of labeled cells in the tissue samples (Fig. 5). The pairwise comparisons reveal distinct performance hierarchies among the viral vectors. FuG-B2 produced significantly higher cell labeling density than AAV2.retro (difference = 166.2, posterior probability = 99.9%, p = 0.003). FuG-B2 also demonstrated superior performance to FuG-E with moderate consistency (difference = 147.7, posterior probability = 99.8%, p = 0.0008) and to FuG-C (difference = 177.7, posterior probability = 99.95%, p < 0.001). Taking into consideration both normalized cell counts in SN (Fig. 4A) and the model-estimated differences accounting for hierarchical structure and individual variability (Fig. 5), the retrograde transduction efficiency of the viral vectors can be ranked as follows: FuG-B2 > FuG-C > FuG-E > AAV2.retro. In our hands, convection enhanced delivery of the same capsid with a different transgene, AAV2.retro-hSyn-CRE-eGFP (NHPs G and H), to the NAc drove expression in the midbrain structure adjacent to SNc, the ventral tegmental area (VTA) (Fig. S2). The midbrain cells transduced in these two cases were not quantified or used for analysis, and dopaminergic identity would need to be validated with a stain against TH.

**Figure 3.**
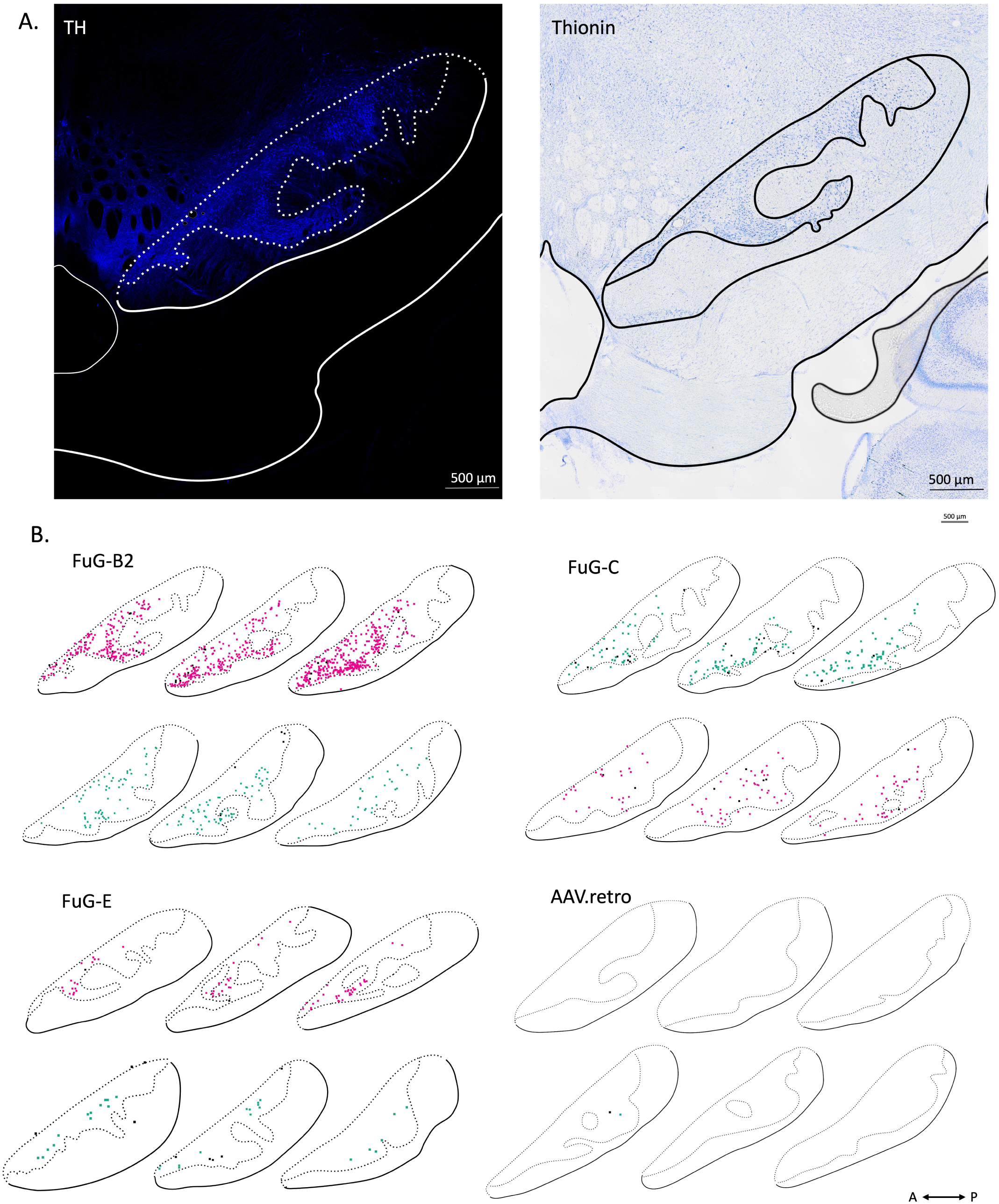
Retrograde expression within SNc. (A) SNc boundary (white dashed line) was determined by the distribution of TH-positive cells (left) and was validated by thionin IHC of a section 160 µm rostral (right). Scale bar 500 µm. (B) Distribution of reporter-positive cells across three consecutive anterior-posterior levels. Cells co-expressing reporter and TH antibody are plotted in color (RFP or GFP), and cells expressing reporter but not TH antibody are plotted in black. Some sections are reflected across the vertical axis for visual symmetry. Data collected from 6 monkeys: FuG-B2 - Monkey A (top), Monkey B (bottom); FuG-C - Monkey A (top), Monkey B (bottom); FuG-E - Monkey C (top), Monkey D (bottom); AAV2.retro - Monkey F (top), Monkey E (bottom). Scale bar 500 µm.

**Figure 4.**
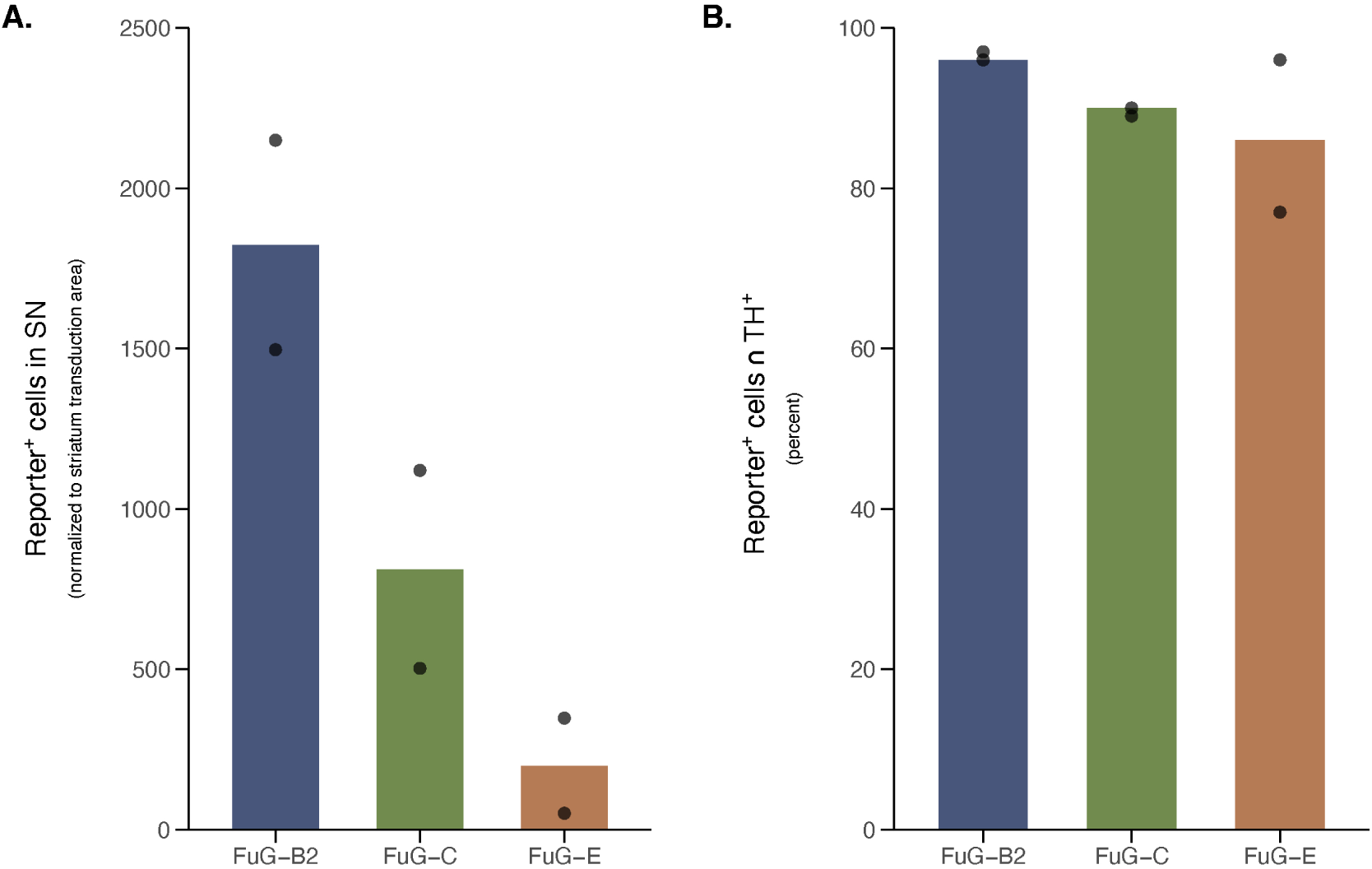
Viral vector expression and specificity. Data points show individual cases per treatment group. (A) Reporter-positive cells in SNc per unit of striatal area transduced (normalization is to cells per 1000 pixels of transduced striatal area). (B) Viral vector selectivity for TH-positive cells within SNc (i.e., of the reporter-positive cells, what percentage of the transduced cells were also TH-positive).

**Figure 5.**
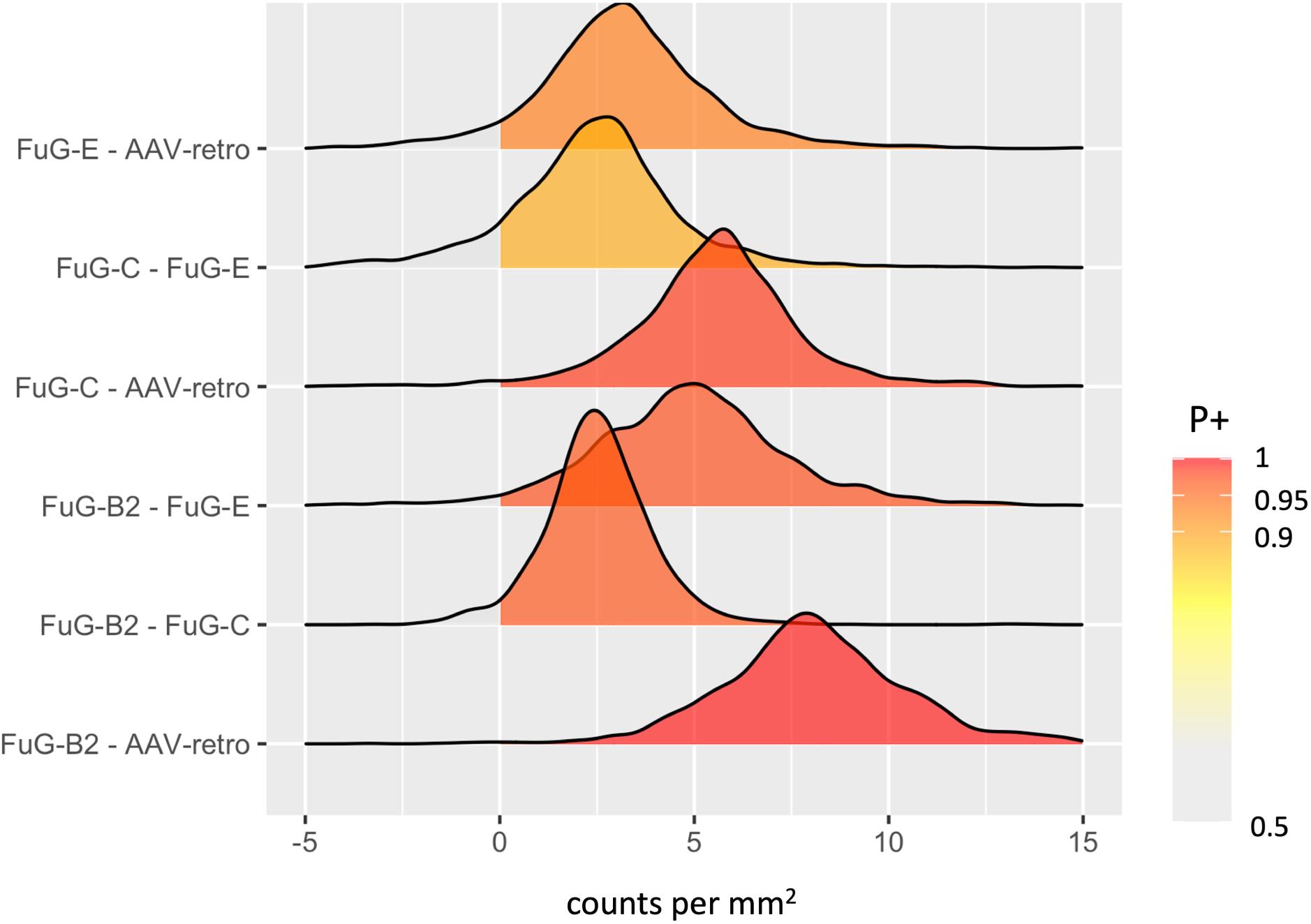
Bayesian modeling. Data was fitted in a hierarchical model under the Bayesian framework, representing different levels of positive effects between treatment groups. Input data indicates reporter^+^ cell counts per unit area on a slice (mm^2^) for each monkey assigned to a treatment group (n = 2 per treatment). The largest positive and consistent differences between treatment groups are indicated in red.

**Figure 6.**
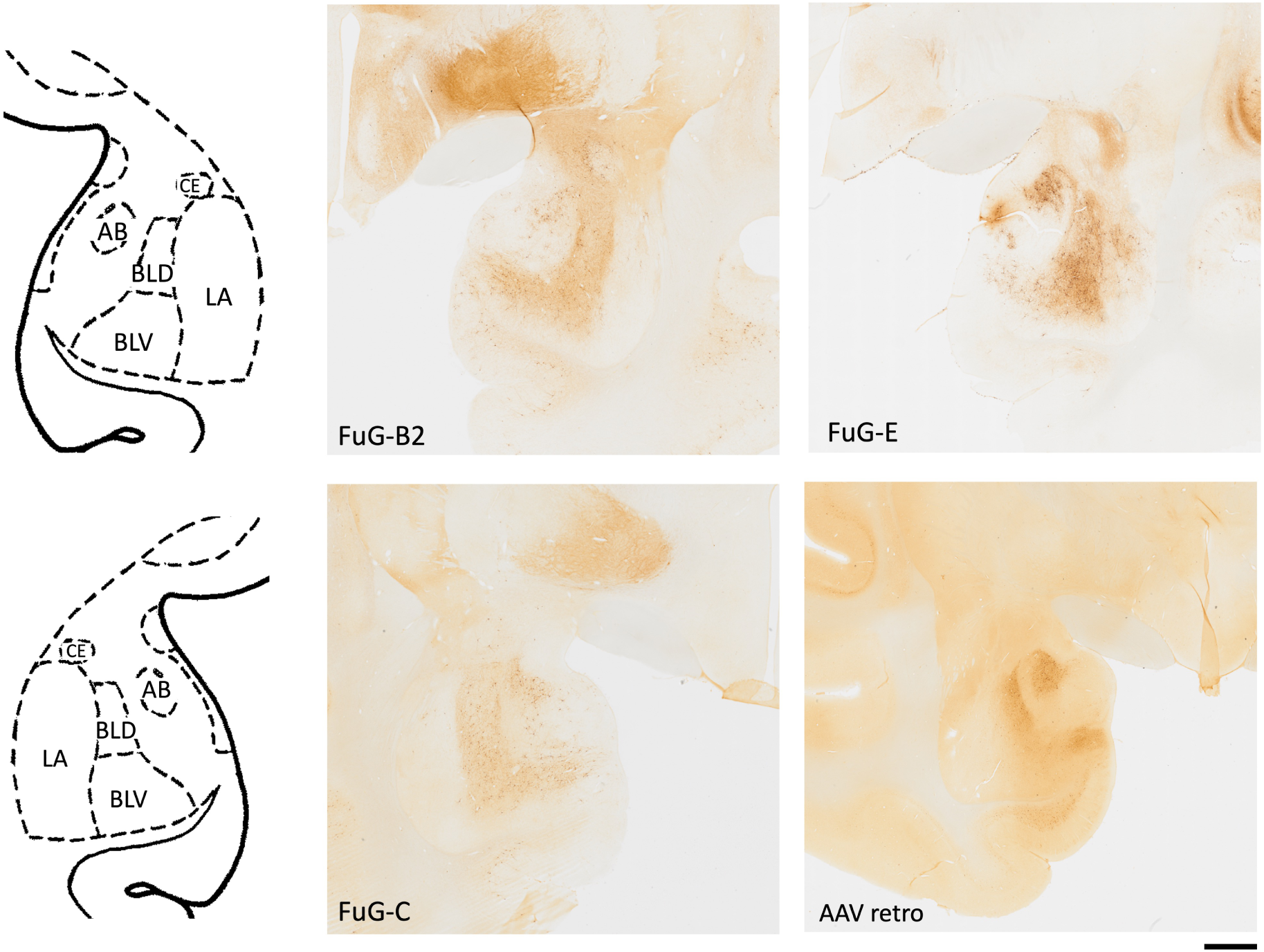
Transduced cells in the BLA following viral delivery into striatum. Schematic of amygdalar subdivisions (top – right hemisphere, bottom – left hemisphere). Retrograde expression within basolateral amygdala. Post-mortem DAB IHC visualized with brightfield microscopy. Scale bar 2 mm. AB – accessory basal nucleus; BLD – basolateral amygdala nucleus, dorsal; BLV – basolateral amygdala nucleus, ventral; LA – lateral amygdala; CE – Central nucleus.

### 3.2 Specificity of viral vector transduction in SNc

FuG-C and FuG-E pseudotypes demonstrate predominantly neuronal tropism, whereas FuG-B2 facilitates gene delivery to neurons and glial cells at the site of injection (Kato & Kobayashi, 2020; Tanabe et al., 2019). Fluorescent reporter sequences (RFP or GFP) in the transfer plasmid were expressed under the control of an hSyn promoter, which confers preferential transduction of neuronal cells, and thereby restricts expression to neurons in the FuG-B2 cases. FuG-B2 exhibited superior selectivity for TH-positive cells in the SNc (96.5%) as evidenced by co-localization of TH-antibody and reporter (Figs. 2B, 3B, 4B), suggesting an intrinsic tropism for dopaminergic neuronal populations. FuG-C also exhibited strong DA cell specificity (89.5%), but transduced other cell types, potentially GABAergic populations within SNc, at a higher rate than FuG-B2. FuG-E showed comparable DA-cell specificity (86.5%) to FuG-C (Fig. 4, Table 2).

### 3.3 Expression and transduction outside of SNc

Although the pseudotyped FuG-B2 vector demonstrated superior efficiency and selectivity within the midbrain following intrastriatal injection, the other three vectors exhibited high transduction in different brain regions, such as cortex and basolateral amygdala (BLA).

AAV2.retro and FuG-E demonstrated qualitatively comparable transduction levels of cells in the basolateral amygdala (BLA) to FuG-B2 and FuG-C (Fig. 6). This contrasts with the failure of AAV2.retro to transduce cells in the SNc, and with the weak FuG-E transduction in the midbrain. Two further NHPs (G and H) received convection-enhanced delivery of the same AAV2.retro capsid with a different transgene, AAV2.retro-hSyn-CRE-eGFP, into the NAc core and shell.

Expression in SN and VTA was evaluated qualitatively (Fig. S2), and this virus appears to transduce cells in the midbrain effectively.

## 4 Discussion

The present study investigated retrograde transduction efficiency and selectivity for dopaminergic cells in the SNc of the NHP using four different viral vectors, FuG-B2, FuG-C, FuG-E, and AAV2.retro. Combining a pan-neuronal promoter, human synapsin (hSyn), with FuG-B2 appeared to improve expression in neuronal populations relative to FuG-C and FuG-E.

### 4.1 Viral vector safety

Viral infection of glial cells can trigger pro-inflammatory response, with microglia (derived from myeloid precursors) and astrocytes playing central roles in neuroinflammation, potentially affecting their supportive functions in neuronal survival and increasing the risk of tumorigenesis (Kobayashi et al., 2017; Rosario et al., 2016; Tanabe et al., 2019). Introducing an hSyn promoter to the construct reduces glial cell transduction at the site of local injection, reducing the risk of a pro-inflammatory response typically seen with transduction of non-neuronal cells (Ciesielska et al., 2013; Samaranch et al., 2014). Contrary to previous findings in which a ubiquitous promoter was used (Tanabe et al., 2019), no gross changes in anatomical integrity were observed in this study following the delivery of the four test vectors, although more sensitive measures of inflammation would need to be performed before using any of these vectors for clinical applications.

### 4.2 AAV2.retro labeling inconsistencies in the midbrain

A recent study (Yan et al., 2025) demonstrated successful retrograde transduction of DA populations in the NHP SNc following intrastriatal delivery of AAV2.retro, which failed to transduce DA cells in our hands. There are several factors which may have explained the discrepancy between Yan’s finding and ours, including the use of a 2A linker (Lewis et al., 2015) and a nuclear localization signal (NLS) (Earley et al., 2014) in the present study. However, a previous study (Cushnie et al., 2020) used AAV2.retro without an NLS or a 2A linker and found dense terminal field labeling, but not labeling of cells in the SNc, ruling out these factors as the explanatory variable for the discrepancies between Yan’s study and ours. However, factors beyond capsid-promoter interactions (Bohlen et al., 2020) (titer, brain area, injection method, species) may also dictate transduction patterns in the NHP, emphasizing the importance of construct composition and methods for replication of findings.

Convection enhanced delivery of the same AAV2.retro capsid with a different transgene, AAV2.retro-hSyn-CRE-eGFP (Fig. S2), successfully drove expression in the midbrain, specifically in the VTA in our hands. The transduction of these cell populations may be attributed to the specific targeting of the NAc shell and core, as well as construct composition (NHPs G and H). The identity of the transduced cells needs to be validated with more extensive staining.

Targeting injections to the putamen and caudate with AAV2.retro (NHPs E and F) yielded sparse labeling in the SNc, like Weiss (2020). Weiss used convection-enhanced delivery in the striatum with AAV2.retro, suggesting that the injection method cannot be the attributing factor for successful expression in SNc. Our observation of AAV2.retro (NHPs E and F) expression in amygdala (Fig. S1) and cortex provides evidence that the striatal injections were successful in driving retrograde transduction, but the specific construct AAV2retro-hSyn1-GCAMP6f-P2A-nls-dTomato failed to express in DA cell populations of the SNc. New engineered AAV2 variants (Dzhashiashvili et al., 2025) show promise for successful expression in NHP brain regions not consistently transduced by AAV2.retro, and may prove to be useful alternatives to the FuG-B2 vector pending more extensive evaluation. Consistency of results across macaque species should also be explored further.

### 4.3 Circuit and cell-type specificity

The utility of viral vector systems lies in their potential to deliver genetic tools with cell-type and circuit specificity. Leveraging retrograde transport to achieve efficient labeling of specific neuronal populations enables therapeutic interventions in primate models of neurodegenerative diseases (Y. Chen et al., 2023). Combining retrograde transport with cell-type specific expression allows for precise targeting of neuromodulatory inputs, thus providing a means of selectively altering circuit-level processing. Populations of neurons in the SNc that project to caudate and putamen are largely segregated (Haber et al., 2000, p. 200; Yan et al., 2025). These subpopulations may display differential vulnerability to pathology or play distinct roles within the circuit. Retrograde transduction with cell-type-selective enhancers, such as AiP13863 (Hunker et al., 2025) may be an effective means of targeting subpopulations of neurons in the SNc, ideal for microcircuit dissection at the level of basic research and for a ‘precision medicine’ approach in gene therapy . Our results suggest that FuG-B2 may be an optimal starting point; combining this vector with cell-type-selective enhancers has the potential to provide the selectivity required to exploit molecular genetic tools to their full potential for the study of the dopaminergic system.

## Supplementary information

## Acknowledgments and Funding

We thank Dr. Mary K L Baldwin for advice on microscopy. Anatomical MRI scanning was carried out in the Neurophysiology Imaging Facility Core (NIMH, NINDS, NEI). This work was supported by the Intramural Research Program, National Institutes of Health, Department of Health and Human Services under project numbers: MH002619-28 (B.J.R.) and NS003135 (Z.M.K.), and by a NARSAD Young Investigator Grant from the Brain and Behavior Research Foundation (MAGE). Funding for this work was also provided by Aligning Science Across Parkinson’s (ASAP-020529) to Z.M.K. through the Michael J. Fox Foundation for Parkinson’s Research (MJFF). This research was supported in part by the Intramural Research Program of the National Institutes of Health (NIH). The contributions of the NIH author(s) were made as part of their official duties as NIH federal employees, are in compliance with agency policy requirements, and are considered Works of the United States Government. However, the findings and conclusions presented in this paper are those of the author(s) and do not necessarily reflect the views of the NIH or the U.S. Department of Health and Human Services.

## Author Contributions

ASP drafted the manuscript with guidance from MAGE. All authors reviewed and edited the manuscript. ASP, VDC, and MAGE performed the surgical procedures. ACC performed histology. ASP performed microscopy. ASP & MAGE analyzed data. WL and LS made the lentiviral vectors. GC developed the statistical analysis for this dataset.

Conceptualization: BA, BJR, ZMK, MAGE.

## Data Availability

The raw data reported in the main text are available online through Figshare (DOI: 10.6084/m9.figshare.30898529).

## Supplementary Figures

**Supplemental Figure 1.**
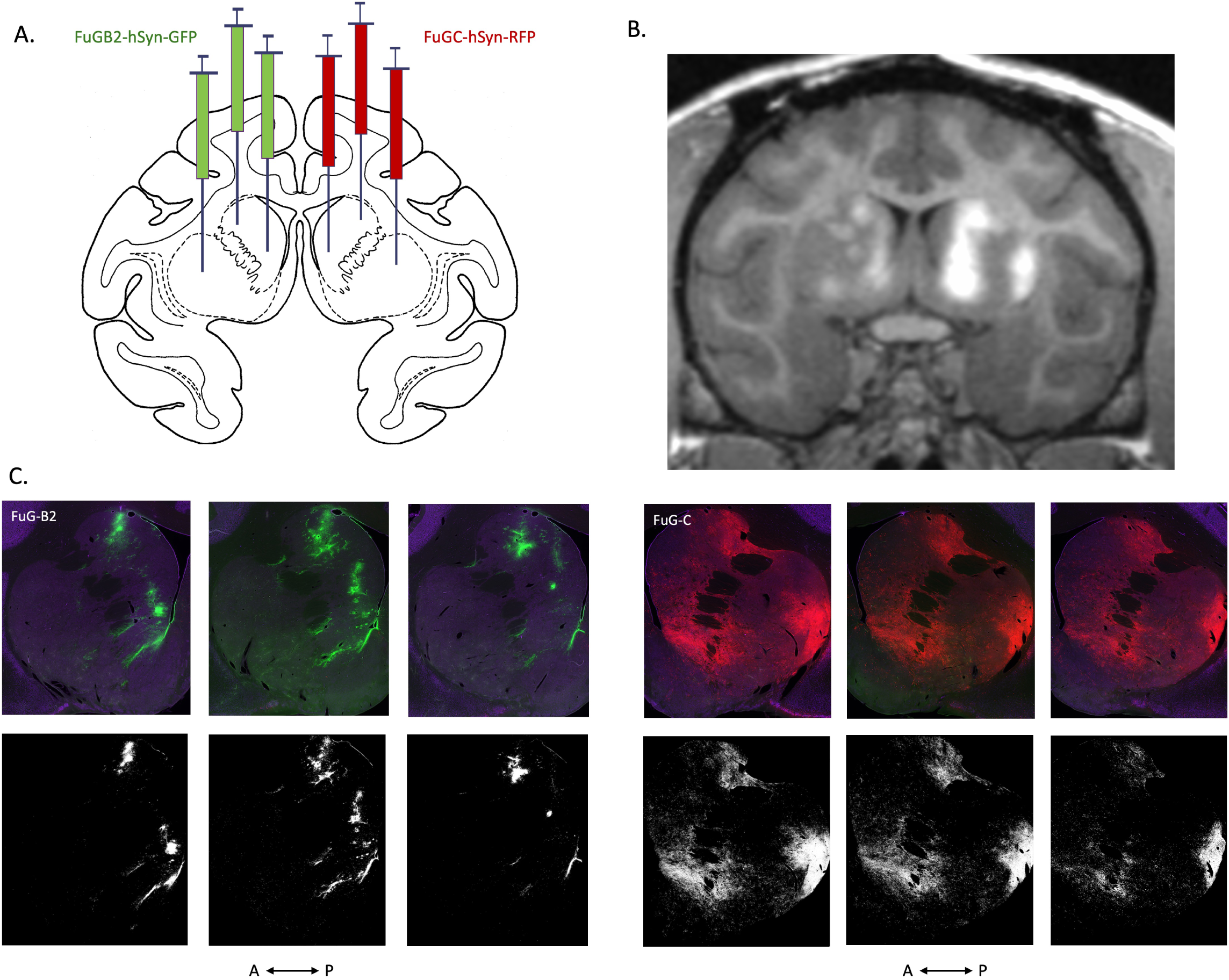
Viral delivery and expression in the striatum of a macaque brain. (A) Schematic illustrates striatal injection sites in Monkey B. FuG-B2 was delivered into the left hemisphere and FuG-C was delivered into the right hemisphere (counter-balanced with Monkey A). (B) Same-day post-op MRI to visualize target accuracy via 0.2 mM manganese contrast agent co-infused with virus. (C) Reporter expression visualized post-mortem with immunohistochemistry and fluorescence microscopy across 3 consecutive anterior-posterior sections with 400 micron spacing. The striatum expression was thresholded using ImageJ for quantification of the injection area (bottom row).

**Supplemental Figure 2.**
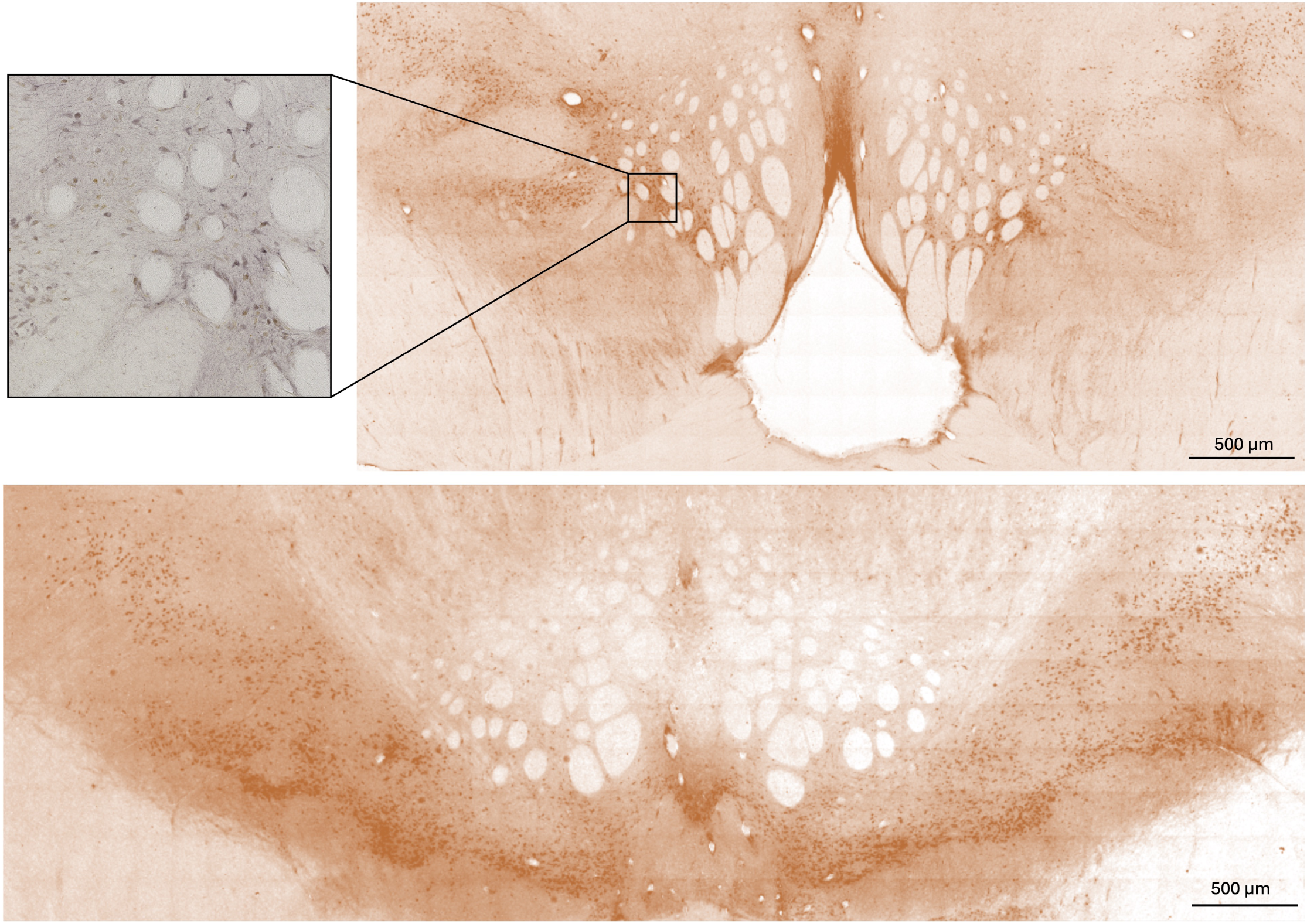
Viral expression in the VTA and SN of a macaque brain following convection enhanced delivery to the NAc (core and shell). 80 µl of AAV2.retro-hSyn-CRE-eGFP (titer 1 x 10^12^ ) was delivered bilaterally to the ventral striatum. Expression visualized 6 weeks post-mortem. Slice thickness 50 µm. Scale bar 500 µm.

